# Hypoxia promotes an inflammatory phenotype of fibroblasts in pancreatic cancer

**DOI:** 10.1101/2022.05.05.490771

**Authors:** Ashley Mello, Tenzin Ngodup, Yusoo Lee, Katelyn L. Donahue, Jinju Li, Arvind Rao, Eileen S. Carpenter, Howard C. Crawford, Marina Pasca di Magliano, Kyoung Eun Lee

## Abstract

Pancreatic ductal adenocarcinoma (PDAC) is characterized by an extensive fibroinflammatory stroma and often experiences conditions of insufficient oxygen availability, or hypoxia. Cancer-associated fibroblasts (CAF) are a predominant and heterogeneous population of stromal cells within the pancreatic tumor microenvironment. Here, we uncover a previously unrecognized role for hypoxia in driving an inflammatory phenotype in PDAC CAFs. We identify hypoxia as a strong inducer of tumor IL1α expression, which is required for inflammatory CAF (iCAF) formation. Notably, iCAFs preferentially reside in hypoxic regions of PDAC. Our data implicate hypoxia as a critical regulator of CAF heterogeneity in PDAC.

## INTRODUCTION

Pancreatic ductal adenocarcinoma (PDAC) remains a deadly disease, with a five-year survival rate of 10% [1]. A notable feature of PDAC is the presence of an abundant fibroinflammatory stroma that includes extracellular matrix (ECM), cancer-associated fibroblasts (CAF), and immune cells [2]. Recently, single-cell RNA-sequencing (scRNA-seq) and other approaches have revealed transcriptionally and functionally distinct CAF subpopulations, myofibroblastic CAFs (myCAF), inflammatory CAFs (iCAF), and antigen-presenting CAFs (apCAF) [3–7]. The myCAF subset is involved in the production of ECM, whereas the iCAF subtype produces high levels of inflammatory cytokines and chemokines [7, 8]. The apCAF population is characterized by MHC class II expression [3]. Previous studies suggested a tumor-restrictive role for myCAFs and a tumor-promoting role for iCAFs and demonstrated that these subpopulations have the potential to interconvert [3, 6–10]. Mechanisms underlying CAF heterogeneity and plasticity as well as different roles of individual CAF subsets in pancreatic tumorigenesis are only beginning to be understood.

Hypoxia, or oxygen (O_2_) deprivation, occurs in solid tumors including PDAC because of their high oxygen/nutrient demand and aberrant vascularization [11–13]. Tumor hypoxia induces adaptive changes in cancer cells and surrounding stromal cells, and is associated with cancer progression and therapy resistance [14, 15]. Although hypoxia has been shown to promote fibrosis and angiogenesis by stimulating fibroblasts [16–18], the relationship between hypoxia and the recently defined CAF subsets in PDAC is unknown.

Here, we show that iCAFs are preferentially located in hypoxic regions of mouse PDAC *in vivo*, and that the hypoxia-related gene signature is positively enriched in iCAFs in human PDAC samples. Using three-dimensional (3D) cocultures of pancreatic cancer cells and fibroblasts, we demonstrate that hypoxia promotes an iCAF state. Our study identifies hypoxia as a key environmental cue for inducing an iCAF phenotype, thus highlighting an instructive role of hypoxia in shaping the stromal microenvironment.

## RESULTS

### CAF subtype proportions differ between normoxic and hypoxic tumor microenvironments

We and others have shown that there is considerable intratumoral heterogeneity of hypoxia in human and mouse PDAC tumors [19, 20]. To identify cells residing in hypoxic tumor areas *in vivo*, we injected Hypoxyprobe, an indicator of pO_2_ levels ≤ 1% [21], intraperitoneally into mice bearing orthotopic PDAC. In this model, pancreatic cancer cells derived from the *Kras^LSL-G12D/+^;Trp53^LSL-R172H/+^;Pdx1-Cre* (KPC) mouse model of PDAC [22] were injected into the pancreas of syngeneic C57/BL6 mice. Immunofluorescence staining for Hypoxyprobe in orthotopic PDAC showed patchy patterns of hypoxia (**Figure 1A**), similar to those observed in human PDAC samples [19]. The average percentage of hypoxic cells in pancreatic tumors (defined as % Hypoxyprobe^+^ cells of total live cells) was 28% (**Figure 1B**). One-third of total PDPN^+^ CAFs stained positively for Hypoxyprobe (**Figures 1C** and **1D**). Using a previously validated flow cytometry strategy for CAF subtypes [3, 6], we evaluated myCAFs (PDPN^+^Ly6C^-^MHCII^-^), iCAFs (PDPN^+^Ly6C^+^MHCII^-^), and apCAFs (PDPN^+^Ly6C^-^MHCII^+^) located within either hypoxic (Hypoxyprobe^+^) or normoxic (Hypoxyprobe^-^) tumor regions (**Figure 1E**). Importantly, the distributions of CAF subpopulations from normoxic and hypoxic tumor microenvironments significantly differed (**Figures 1E-1I** and **S1A**). Although myCAFs were the prevalent CAF subset in both normoxic and hypoxic tumor regions (**Figures 1F and 1I**), hypoxic areas contained a significantly higher fraction of iCAFs compared with normoxic areas (**Figures 1G and 1I**) and exhibited pronounced increases in the iCAF/myCAF ratio (**Figure 1H**). We further confirmed the preferential localization of iCAFs in hypoxic regions in orthotopic tumors using a different PDAC cell line, 4662 (**Figure S1B**).

**Figure 1.**
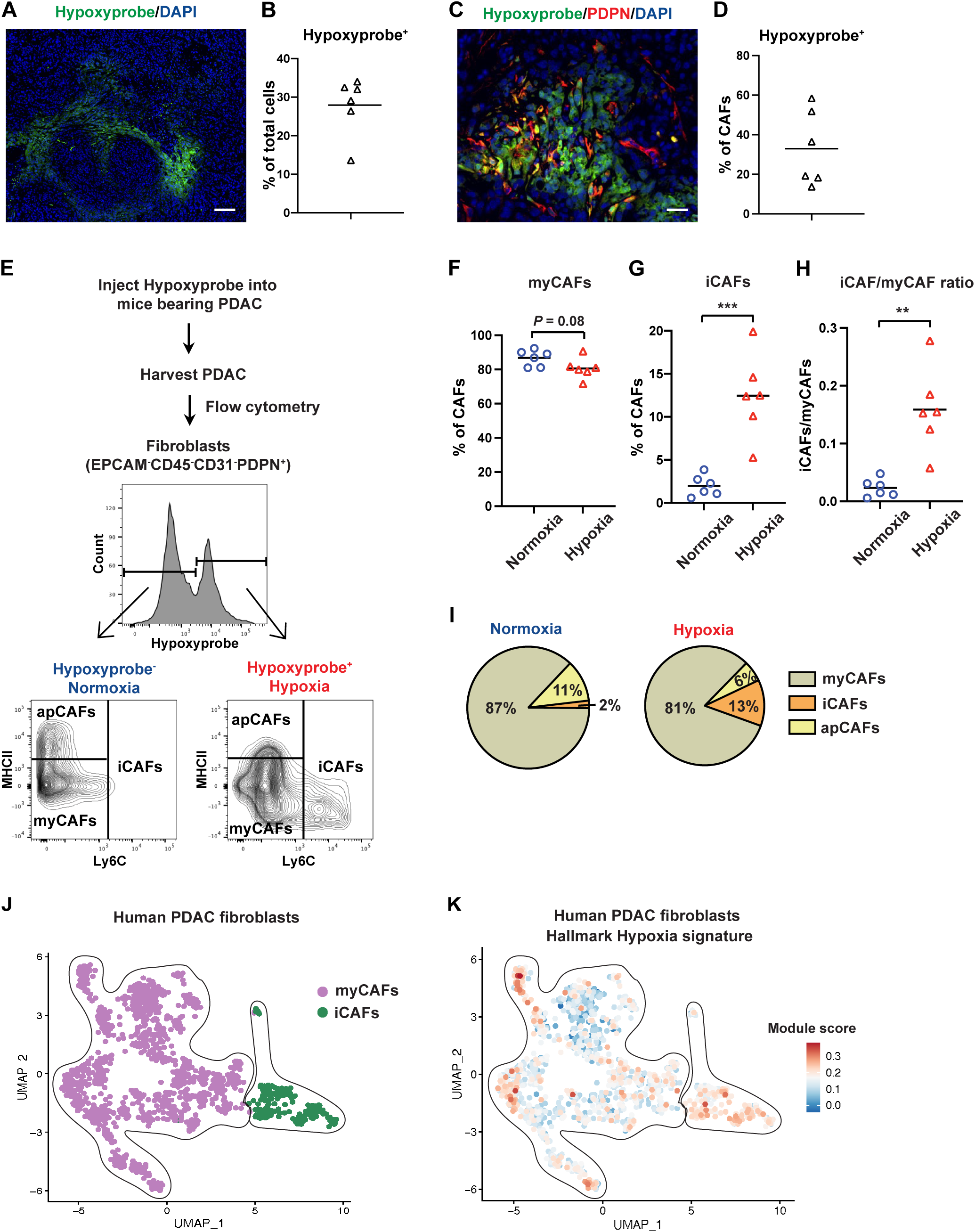
iCAFs preferentially reside in hypoxic regions of PDAC *in vivo*. (A-I) Mice bearing 4-week orthotopic PDAC of mT3 tumor cells received an intraperitoneal injection with 60 mg/kg of Hypoxyprobe and were sacrificed 1.5–2 hours later. (A) Immunofluorescence staining for Hypoxyprobe (green) and DAPI (blue) in orthotopic PDAC. Scale bar, 100 μm. (B) Percentage of Hypoxyprobe^+^ cells among total live cells from orthotopic PDAC, as analyzed by flow cytometry (n=6). (C) Co-immunofluorescence staining for Hypoxyprobe (green), PDPN (red), and DAPI (blue) in orthotopic PDAC. Scale bar, 25 μm. (D) Percentage of Hypoxyprobe^+^ CAFs among total CAFs from orthotopic PDAC, as analyzed by flow cytometry (n=6). (E) Schematic of flow cytometry strategy to identify CAF subsets residing in normoxic and hypoxic tumor regions. Representative flow plots showing the gating strategy for the analysis of normoxic (Hypoxyprobe^-^) and hypoxic (Hypoxyprobe^+^) CAF subsets from orthotopic PDAC. (F-H) Percentage of myCAFs (F), percentage of iCAFs (G), and iCAF/myCAF ratio (H) among normoxic and hypoxic CAFs from orthotopic PDAC, as analyzed by flow cytometry (n=6). (I) Pie charts showing mean frequencies of the indicated subsets among normoxic and hypoxic CAFs from orthotopic PDAC, as analyzed by flow cytometry (n=6). (J) Uniform and manifold approximation and projection (UMAP) visualization of fibroblast clusters from human PDAC scRNA-seq (n=16 patients merged). Different CAF subtype clusters are color-coded. Data are from Steele and colleagues [23], and annotations are from Kemp and colleagues [24]. (K) UMAP visualization of human PDAC fibroblasts from (J) colored by hypoxia gene set expression score. The hypoxia signature for analysis was obtained from MSigDB’s Hallmark collection. Red, highest score of hypoxia signature; blue, lowest score of hypoxia signature. The symbols in (B, D, F, G, and H) represent individual mice, and horizontal lines represent the means. P values were determined by student’s *t* test. **p < 0.01; ***p < 0.001.

To address the relevance of correlation between hypoxia and an iCAF phenotype in human PDAC, we interrogated the expression profiles of iCAFs and myCAFs from a scRNA-seq data set [23] that includes 16 PDAC patient tumor samples. Populations of myCAFs and iCAFs were both present in this dataset, with the majority of fibroblasts falling into the myCAF group (**Figure 1J,** as annotated in [24]). The expression profiles of each cell were then scored using the Hallmark Hypoxia gene set (MSigDB) as a readout of exposure to hypoxia. We found that most iCAFs exhibit a robust hypoxia profile (76%, 177 of 234 total cells scored above the median of signature expression), while only a subset of myCAFs met this threshold (45%, 565 of 1,250 total cells scored above the median of signature expression) (**Figure 1K**). These observations suggest that the iCAF phenotype is linked with the hypoxic tumor microenvironment of PDAC.

### Hypoxia promotes the induction of an inflammatory phenotype in CAFs by modulating their interactions with tumor cells

Based on the correlation between PDAC hypoxia and iCAF enrichment, we set out to determine whether hypoxia regulates an iCAF phenotype. When pancreatic stellate cells (PSCs), a precursor population of CAFs, are seeded in Matrigel in a transwell insert and cultured with PDAC tumor organoids in the lower compartment of the plate, they acquire the inflammatory features characteristic of iCAFs [7, 8]. On the other hand, PSCs cultured alone in Matrigel maintain a quiescent state [7, 8]. To examine the effects of hypoxia on an iCAF phenotype, we exposed the cocultures of mouse PDAC tumor organoids and PSCs to either normoxia (21% O_2_) or hypoxia (1% O_2_) (**Figure 2A**) and measured expression of CAF subset markers in PSCs. Expression of the iCAF markers *Il6, Cxcl1*, and *Lif* was markedly elevated in hypoxic PSCs cocultured with tumor organoids relative to their normoxic counterparts (**Figures 2B** and **2C**). On the other hand, expression of the myCAF markers *Acta2* (α-SMA gene) and *Transgelin* (*Tagln*) in PSCs cocultured with tumor organoids was not affected by hypoxia (**Figure 2D**). Importantly, hypoxic induction of the iCAF markers in PSCs only occurred when cocultured with tumor organoids but not when cultured alone (**Figure 2B**). These observations were reproduced in the cocultures of tumor cells and a CAF line derived from mouse PDAC (**Figure S2**), suggesting that hypoxic induction of the iCAF phenotype is not limited to PSC-derived CAFs. To assess the effects of hypoxia on fibroblast phenotype in an unbiased fashion, we performed RNA-seq profiling of the hypoxic and normoxic PSCs cocultured with PDAC organoids. When using gene-set enrichment analysis (GSEA), we found that “inflammatory response” and “IL6/JAK/STAT3”, in addition to the “hypoxia signature” are top-ranked in association with the hypoxic PSCs (**Figure 2E**). Altogether, these results indicate that hypoxia promotes an iCAF state and that the induction of an inflammatory fibroblast phenotype by hypoxia requires factors secreted by tumor cells.

**Figure 2.**
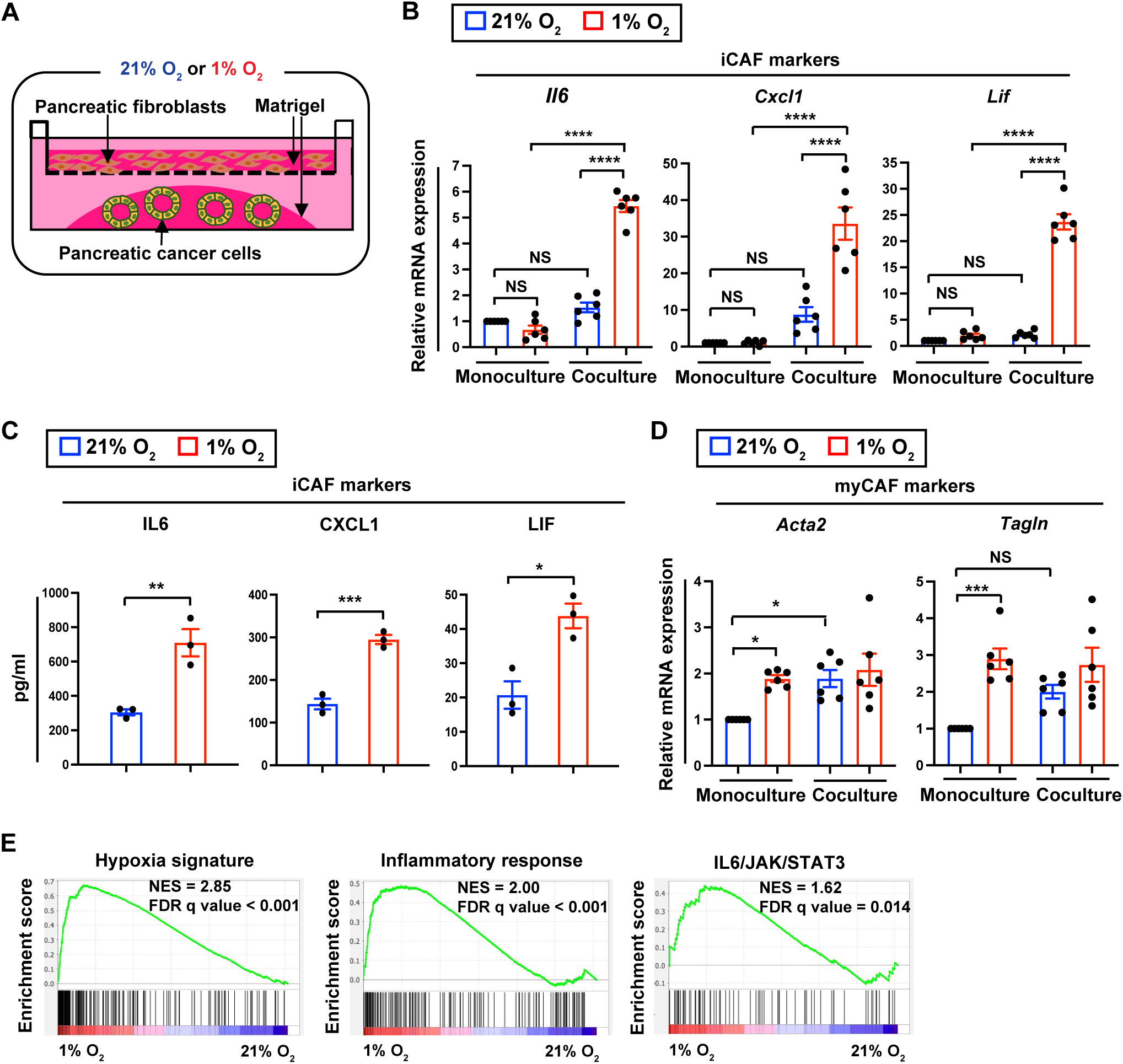
Hypoxia promotes an inflammatory fibroblast phenotype. (A) Schematic illustration of the 3D coculture platform to model tumor cell-fibroblast interactions under normoxia and hypoxia. (B) Quantitative RT-PCR analysis of iCAF markers in PSCs cultured alone or with mT3 tumor organoids under 21% O_2_ or 1% O_2_ for 48 hours (n=6). Expression levels were normalized by *18S rRNA*. (C) Enzyme-linked immunosorbent assay (ELISA) of iCAF markers in conditioned media from 3D cocultures of PSCs and mT3 tumor cells under 21% O_2_ or 1% O_2_ for 72 hours (n=3). (D) Quantitative RT-PCR analysis of myCAF markers in PSCs cultured alone or with mT3 tumor organoids under 21% O_2_ or 1% O_2_ for 48 hours (n=6). Expression levels were normalized by *18S rRNA*. (E) Gene-set enrichment analysis (GSEA) showing significantly upregulated pathways in PSCs cultured with mT3 tumor organoids at 1% O_2_ compared with PSCs cultured with mT3 tumor organoids at 21% O_2_ for 48 hours. NES, normalized enrichment score; FDR, false discovery rate. Each data point in (B-D) represents individual primary PSC lines. Data in (B-D), mean±SEM. P values were determined by two-way ANOVA with Bonferroni post-test (B and D) and student’s *t* test (C). NS, not significant. *p < 0.05; **p < 0.01; ***p < 0.001; ****p < 0.0001.

### Hypoxic regulation of the iCAF phenotype is independent of tumor HIF1α or HIF2α

Cellular adaptation to hypoxia is largely coordinated by hypoxia-inducible factors (HIF) [25]. It has been shown that the major HIF isoforms, HIF1α and HIF2α, are expressed in human and mouse PDAC and play distinct roles in pancreatic tumorigenesis [20, 26–28]. Because hypoxic tumor cells are needed to establish the iCAF phenotype (**Figure 2B**), we postulated that HIF1α or HIF2α in tumor cells may contribute to iCAF formation under hypoxia. To test this hypothesis, we knocked down HIF1α or HIF2α in PDAC tumor cells using shRNAs (**Figures S3A** and **S3B**) and cultured these tumor cells with PSCs under normoxia (21% O_2_) or hypoxia (1% O_2_). Unexpectedly, neither HIF1α knockdown nor HIF2α knockdown impaired induction of iCAF marker genes *Lif* and *Cxcl1* in PSCs by hypoxia (**Figures 3A** and **3B**). These data suggest that tumor HIF1α and HIF2α are dispensable for hypoxia-mediated iCAF formation.

**Figure 3.**
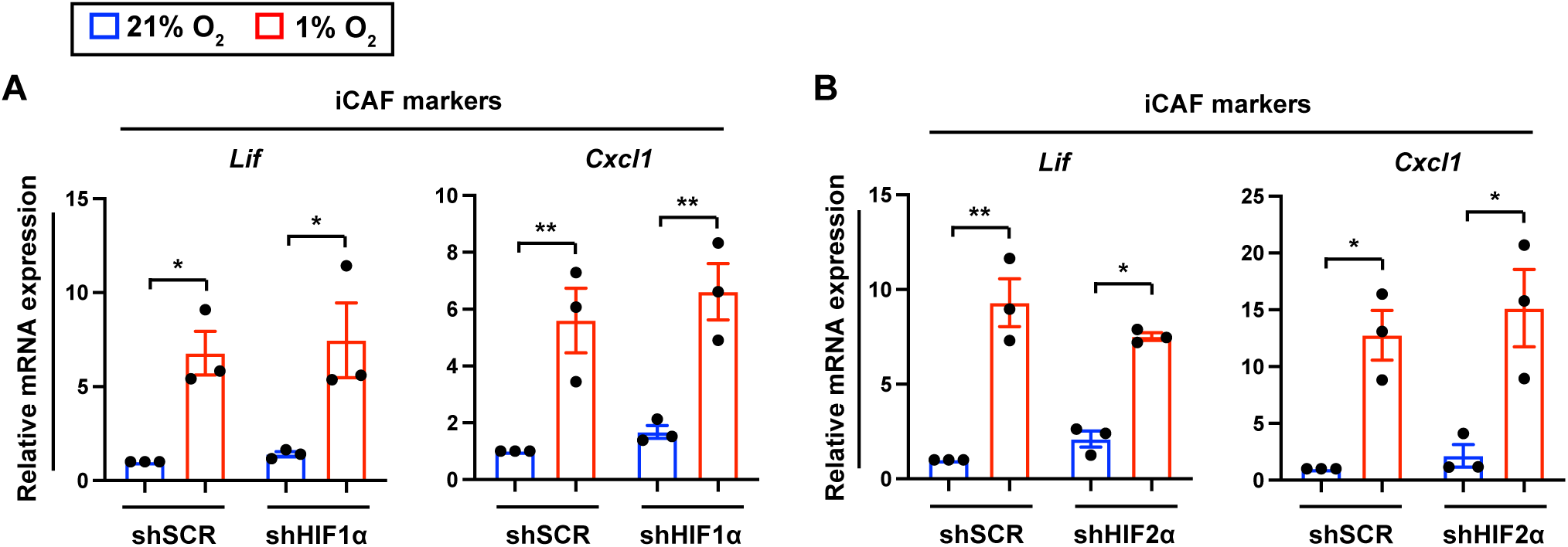
Hypoxia induces inflammatory fibroblasts in a tumor HIF-independent manner. (A and B) Quantitative RT-PCR analysis of iCAF markers in PSCs cultured with mT3 tumor organoids expressing scrambled shRNA (shSCR) control, HIF1α shRNA (shHIF1α) (A), or HIF2α shRNA (shHIF2α) (B) under 21% O_2_ or 1% O_2_ for 48 hours (n=3). Expression levels were normalized by *18S rRNA*. Each data point represents individual primary PSC lines. Results show mean±SEM. P values were determined by two-way ANOVA with Bonferroni post-test. *p < 0.05; **p < 0.01.

### IL1α in tumor cells mediates hypoxic induction of the iCAF phenotype

IL1α secreted by pancreatic tumor cells and subsequent IL6/JAK/STAT3 activation in CAFs have been shown to trigger iCAF formation [7]. However, the mechanism underlying IL1α induction in cancer cells has remained obscure. Because iCAF induction by hypoxia requires tumor cells, we measured IL1α expression from tumor organoids exposed to either normoxia (21% O_2_) or hypoxia (1% O_2_). Hypoxia increased *Illa* mRNA levels in pancreatic cancer cells (**Figure 4A**). IL1α protein levels were also elevated in conditioned media from hypoxic cocultures relative to conditioned media from normoxic cocultures (**Figure 4B**). Of note, although hypoxia significantly upregulated *Illa* expression in cancer cells cultured alone, an increase in *Illa* expression in cancer cells by hypoxia was even greater in the presence of PSCs (**Figure 4A**), implicating bi-directional interactions between tumor cells and PSCs. Moreover, targeting IL1α with a neutralizing antibody substantially reduced induction of iCAF marker genes *Lif* and *Cxcl1* in PSCs but not myCAF marker gene *Acta2* under hypoxia (**Figures 4C** and **S4A**). To confirm the importance of IL1 signaling in iCAF formation, we treated cancer cell-PSC cocultures with an IL1 receptor (IL1R1)-neutralizing antibody, which resulted in the impaired acquisition of the iCAF phenotype under hypoxia (**Figure 4D**). Consistent with a tumor HIF-independent mechanism for the hypoxic induction of iCAF formation, knockdown of HIF1α or HIF2α did not affect upregulation of *Illa* in cancer cells in response to hypoxia (**Figures S4B** and **S4C**). Collectively, our findings suggest that hypoxia induces IL1α expression in tumor cells and that IL1α is critical for hypoxia-mediated iCAF formation.

**Figure 4.**
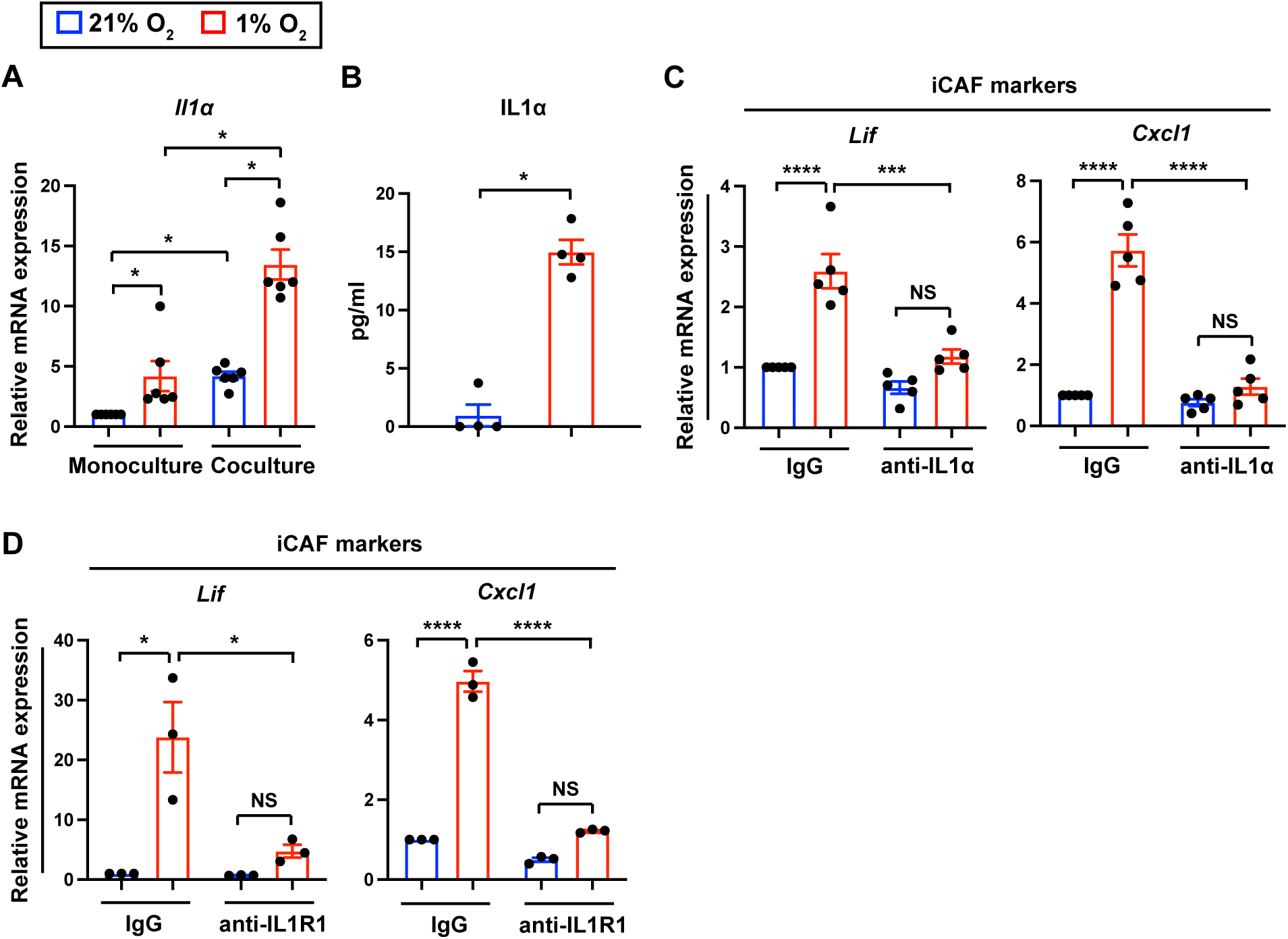
IL1α in tumor cells mediates the induction of the iCAF phenotype under hypoxia. (A) Quantitative RT-PCR analysis of *Il1a* in mT3 tumor organoids cultured alone or with PSCs under 21% O_2_ or 1% O_2_ for 48 hours (n=6). Expression levels were normalized by *18S rRNA*. (B) ELISA of IL1α in conditioned media from 3D cocultures of mT3 tumor cells and PSCs under 21% O_2_ or 1% O_2_ for 72 hours (n=4). (C) Quantitative RT-PCR analysis of iCAF markers in PSCs cultured with mT3 tumor organoids in the presence of IL1α-neutralizing antibody or isotype control antibody under 21% O_2_ or 1% O_2_ for 72 hours (n=5). Expression levels were normalized by *18S rRNA*. (D) Quantitative RT-PCR analysis of iCAF markers in PSCs cultured with mT3 tumor organoids in the presence of IL1R1-neutralizing antibody or isotype control antibody under 21% O_2_ or 1% O_2_ for 72 hours (n=3). Expression levels were normalized by *18S rRNA*. Each data point in (A-D) represents individual primary PSC lines. Data in (A-D), mean±SEM. P values were determined by Mann-Whitney test with Bonferroni post-test (A), Mann-Whitney test (B), and two-way ANOVA with Bonferroni post-test (C and D). NS, not significant. *p < 0.05; ***p < 0.001; ****p < 0.0001.

## DISCUSSION

Hypoxia is a critical feature of the tumor microenvironment and predicts poor clinical outcome [14, 15]. The impact of hypoxia on cancer cells has been well-characterized, yet much remains to be understood as to how hypoxia regulates stromal components and the tumor-stroma crosstalk. In this study, we demonstrate that intratumoral normoxic and hypoxic microenvironments differ in CAF composition in mouse PDAC and that iCAFs are linked to tumor hypoxia in human and mouse PDAC. By exposing 3D cocultures of pancreatic cancer cells and fibroblasts to either hypoxia or normoxia, we found that hypoxia promotes an inflammatory phenotype of fibroblasts. In addition, we showed that hypoxic induction of the iCAF phenotype requires IL1α emanating from tumor cells and the presence of hypoxic fibroblasts further elevates IL1α levels in tumor cells, implicating hypoxia as a modulator of reciprocal interactions between cancer cells and fibroblasts. At present, the precise mechanism for tumor IL1α regulation by hypoxia remains unknown.

PSCs have been thought to give rise to the majority of PDAC CAFs. However, recent reports suggest that besides PSCs, PDAC CAFs can arise from multiple cell types [29, 30]. It is unclear whether CAF populations of different developmental origins have differential capacity to gain iCAF features. We observed that hypoxia propels an inflammatory phenotype in a PDAC CAF line as well as primary PSCs, suggesting that hypoxia-mediated iCAF induction is not limited to PSCs.

A key finding in our current study is that iCAFs are enriched in hypoxic zones of PDAC compared with normoxic tumor regions. This spatial link between hypoxia and iCAF enrichment *in vivo* together with our *in vitro* finding that hypoxia promotes iCAF induction, raises the possibility that hypoxia plays an active role in driving regional stromal heterogeneity. Notably, recent studies have observed a correlation between iCAF enrichment and immunosuppression [6, 10, 31], which warrants the investigation of the effects of hypoxia on CAF-immune crosstalk.

In summary, our study reveals that the normoxic and hypoxic microenvironments of PDAC exhibit distinct CAF compositions. We also show that hypoxia induces an inflammatory fibroblast phenotype through upregulation of tumor IL1α, thus highlighting the significance of hypoxia in shaping the tumor stroma. Better understanding of the impact of hypoxia on CAF heterogeneity and function is needed to make stroma-targeting therapies clinically viable.

## MATERIALS AND METHODS

### Mice

All animal protocols were reviewed and approved by the Institutional Animal Care and Use Committee of the University of Michigan. Wild-type (WT) C57/BL6 mice (stock # 000664) from Jackson Laboratory were used for PSC isolation and orthotopic transplantation experiments at 8-12 weeks of age, including male and female mice. For orthotopic transplantation, 7.5 × 10^4^ mT3 (provided by Dr. David A. Tuveson) [32] or 7.5 × 10^4^ 4662 cells (provided by Robert H. Vonderheide) [33] derived from primary PDAC in *Kras^LSL-G12D/+^; Trp53^LSL-R172H/+^; Pdx1-Cre* mice of a C57/BL6 genetic background, were resuspended as a 30 μl suspension of 50% Matrigel (#356231, Corning) in PBS and injected into the pancreas. At 4 weeks post transplantation, mice received an intraperitoneal injection with 60 mg/kg of Hypoxyprobe (pimonidazole hydrochloride, Hypoxyprobe, Inc) and were sacrificed 1.5–2 hours later for flow cytometry analysis.

### Cell Lines and Cell Culture

PSCs were isolated from WT mice by enzymatic digestion of pancreatic tissue and subsequent density gradient centrifugation as previously described [8, 34]. Primary PSC lines between passage 2 and 4 were used for all experiments. The FB1 CAF line was generated from an iKras* p53* mouse [35] by fluorescence-activated cell sorting of PDGFRα^+^;EPCAM^-^ cells. The mT3 (provided by Dr. David A. Tuveson) [32] and 4662 (provided by Robert H. Vonderheide) [33] PDAC cell lines were derived from primary murine PDAC. FB1, mT3, 4662 cell lines were cultured no more than 20 to 25 passages. All cells were passaged in DMEM with 10% FBS and 1% penicillin/streptomycin (Thermo Fisher). For 3D cocultures, PSCs were seeded in Matrigel (#356231, Corning) in a transwell insert (#662610, Greiner Bio-One) and cultured with PDAC tumor organoids in the lower compartment of the 24-well plate in DMEM containing 5% FBS and 1% penicillin/streptomycin (Thermo Fisher). For IL1α neutralization experiments, cocultures were treated with 3 μg/ml IL1α-neutralizing antibody (#MAB4001, R&D Systems) or isotype control antibody (#400902, BioLegend) for 72 hours. For IL1R1 neutralization experiments, cocultures were treated with 0.5 μg/ml IL1R1-neutralizing antibody (#PA5-47937, Invitrogen) or isotype control antibody (#AB108C, R&D Systems) for 72 hours. Cell line authentication for FB1 and mT3 was not performed. The 4662 cells were authenticated by the Research Animal Diagnostic Laboratory (RADIL) at the University of Missouri. Mycoplasma testing (MycoAlert Detection Kit, Lonza) was performed monthly.

### Lentiviral-mediated shRNA Transduction

The mT3 PDAC cell line was transduced with lentivirus containing shRNA plasmids at optimized viral titers. Stable cell lines were established post-puromycin selection. The following shRNA plasmids were used: pGIPZ Scrambled shRNA (#RH4346, Horizon), pGIPZ HIF1α shRNA (#RMM4431-200404026, Horizon), pLKO.1 Scrambled shRNA (#1864, Addgene), pLKO.1 HIF2α shRNA (#TRCN0000082307, Sigma).

### Quantitative RT-PCR

Total RNA was isolated from cells using the RNeasy mini kit (#74104, Qiagen). cDNA was synthesized using a High-Capacity cDNA Reverse Transcription Kit (#4368814, Applied Biosystems). PCR reactions were performed using SYBR Green PCR reagents (#A25742, Applied Biosystems) mixed with indicated cDNAs and primers (primer sequences are listed in Supplemental Table S1) in a QuantStudio Real-Time PCR system (Applied Biosystems). Expression levels were normalized by *18S rRNA*.

### Immunofluorescence

Tissues were fixed in 4% paraformaldehyde/PBS (4**°**C, overnight) and processed for paraffin-embedding. For immunofluorescence, slides were boiled for 20 min in 10 mM sodium citrate (pH 6.0) for antigen retrieval and blocked with 5% serum/0.3% Triton X-100 for 1 hour. Sections were incubated with FITC-conjugated Hypoxyprobe-1-MAb1 (4.3.11.3, #FITC-Mab, 1:500, Hypoxyprobe, Inc) and Alexa Fluor 594-conjugated PDPN antibody (8.1.1, #127414, 1:250) diluted in 1% BSA/0.3% Triton X-100 overnight at 4**°**C. Slides were counterstained with DAPI (Invitrogen), and mounted in Prolong Gold antifade reagent (Invitrogen). Fluorescence images were acquired using an Olympus IX73 microscope.

### Flow Cytometry

Single cell suspensions from mouse tissues were prepared as previously described [20]. Cells were stained in PBS/0.5% FBS/2mM EDTA with the following fluorochrome-conjugated antibodies: BV421-conjugated anti-Ly6C (HK1.4, #128031, 1:100), PerCP-Cy5.5-conjugated anti-CD45 (30-F11, #103132, 1:200), PE-conjugated anti-EPCAM (G8.8, #118205, 1:200), PE-conjugated anti-CD31 (390, #102407, 1:200), PE-Cy7-conjugated anti-PDPN (8.1.1, #127411, 1:100), APC-Cy7-conjugated anti-MHCII (M5/114.15.2, #107628, 1:300) (from BioLegend); FITC-conjugated Hypoxyprobe-1-MAb1 (4.3.11.3, #FITC-Mab, 1:200) (from Hypoxyprobe, Inc). The viability marker Zombie Aqua was purchased from BioLegend (#423102). Flow cytometry was performed on a ZE5 Cell Analyzer (Bio-Rad), and data was analyzed using FlowJo software.

### ELISA

For ELISA of media, 3D cocultures were grown under 21% O_2_ or 1% O_2_ for 72 hours. Media were collected, spun down, and assayed using the manufacturer’s protocol. ELISA assays used were IL1α (#433404, BioLegend), CXCL1 (#EMCXCL1, Invitrogen), IL6 (#DY406-05, R&D Systems), and LIF (#445104, BioLegend).

### RNA-seq and Data Analysis

Total RNA was isolated from cells using the RNeasy mini kit (#74104, Qiagen). Libraries were constructed using NEB polyA RNA ultra II and subsequently subjected to 150 cycles of sequencing on NovaSeq-6000 (Illumina). Adapters were trimmed using Cutadapt (v2.3). FastQC (v0.11.8) was used to ensure the quality of data. Reads were mapped to the mouse genome (GRCm38) using STAR (v2.6.1b) and assigned count estimates to genes with RSEM (v1.3.1). Alignment options followed ENCODE standards for RNA-seq. FastQC was used in an additional post-alignment step to ensure that only high-quality data were used for expression quantitation and differential expression. Differential gene expression analysis was performed using DESeq2, using a negative binomial generalized linear model (thresholds: linear fold change >1.5 or <-1.5, Benjamini-Hochberg FDR (Padj) <0.05). GSEA was performed using GSEA 4.1.0.

### Single-cell RNA-seq Analysis

Human single-cell RNA-seq (scRNA-seq) data were previously published in [23] and fibroblasts were annotated in [24]. Both raw and processed data are available at the NIH dbGaP database (accession #phs002071.v1.p1;[23]), with full clinical annotation. Downstream analysis was performed using Seurat V4.0.3 [36]. Hypoxia signature scoring was performed using Seurat’s “AddModuleScore” function.

### Statistical Analysis

Data were analyzed using GraphPad Prism 7 software. Statistical tests with normally distributed variables included two-tailed Student’s *t* test and two-way ANOVA. D’Agostino and Pearson test and/or Shapiro–Wilk test was used to test the normality of sample distribution. When variables were not normally distributed, we performed non-parametric Mann-Whitney test. Bonferroni correction was applied for multiple comparisons. *P* value < 0.05 was considered statistically significant.

## ACKNOWLEDGMENTS

The authors are grateful to Dr. D. A. Tuveson for providing the mT3 PDAC cell line. The 4662 PDAC cell line was a generous gift from Dr. R.H. Vonderheide. We acknowledge support from the Flow Cytometry Core, the Advanced Genomics Core, and the Bioinformatics Core of the University of Michigan Medical School’s Biomedical Research Core Facilities. This project was supported by DOD Peer Reviewed Cancer Research Program grant W81XWH-20-1-0629, Concern Foundation Conquer Cancer Now Award, and Swim Across America Young Investigator Award to K.E.L. A.M. was supported by NIH grant T32AI007413. T.N. was supported by NIH grant T32GM140223. K.L.D. was supported by NIH grant T32CA009676. A.R. was funded by NIH grant R37CA214955, the American Cancer Society RSG-005-016, MIDAS PODS funding, and institutional startup fund from the University of Michigan. E.S.C. was funded by American College of Gastroenterology Junior Faculty Development Award and VA Career Development Award BLR&D. H.C.C. was supported by NIH grant R01CA247516. M.P.d.M. was supported by NIH grants R01CA151588 and R01CA198074, and the American Cancer Society. J.L., H.C.C., and M.P.d.M. were supported by the University of Michigan Cancer Center Support Grant P30CA046592. H.C.C. and M.P.d.M. were supported by NIH grant U01CA224145.

## AUTHOR CONTRIBUTIONS

Conceptualization, K.E.L.; Methodology, A.M., T.N., Y.L., K.L.D., E.S.C., H.C.C., M.P.d.M., K.E.L.; Validation, A.M., T.N., K.E.L.; Formal Analysis, A.M., T.N., K.L.D., J.L., E.S.C., K.E.L.; Investigation, A.M., T.N., Y.L., K.E.L.; Resources, K.L.D., A.R., H.C.C., M.P.d.M., K.E.L.; Data Curation, A.M., T.N., K.L.D., J.L., E.S.C., K.E.L.; Visualization, A.M., T.N., K.L.D., J.L., E.S.C., K.E.L.; Writing – Original Draft, K.E.L.; Writing – Review & Editing, A.M., T.N., K.L.D., M.P.d.M., K.E.L.; Funding Acquisition, K.E.L.; Supervision, K.E.L.

## DECLARATION OF INTERESTS

The authors declare no competing interests.

## Notes

**Disclosure of potential conflicts of interest**: The authors disclose no potential conflicts of interest.

### Competing Interest Statement

The authors have declared no competing interest.

