## Supplemental information for "Hypoxia promotes an inflammatory phenotype of fibroblasts in pancreatic cancer"

#### Figure S1

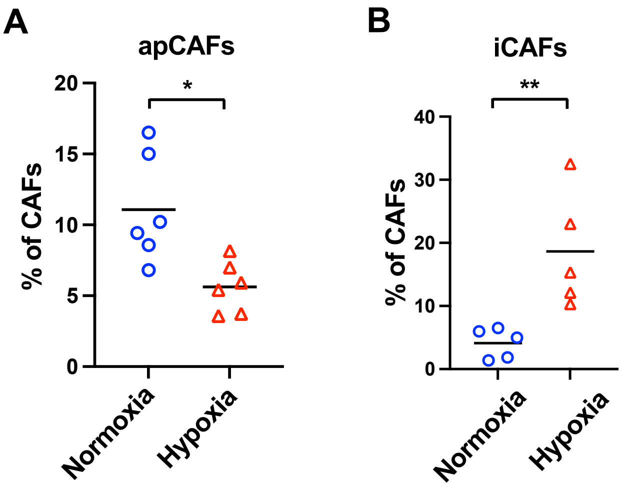


**Figure S1. Distinct CAF subset proportions in normoxic and hypoxic tumor regions.**

Mice bearing 4-week orthotopic PDAC received an intraperitoneal injection with 60 mg/kg of Hypoxyprobe and were sacrificed 1.5–2 hours later.

(A) Percentage of apCAFs among normoxic and hypoxic CAFs from 4-week orthotopic PDAC of mT3 tumor cells, as analyzed by flow cytometry (n=6).

(B) Percentage of iCAFs among normoxic and hypoxic CAFs from 4-week orthotopic PDAC of 4662 tumor cells, as analyzed by flow cytometry (n=5).

The symbols in (A and B) represent individual mice, and horizontal lines represent the means. P values were determined by student's *t* test. *$p < 0.05$; **$p < 0.01$.

#### Figure S2

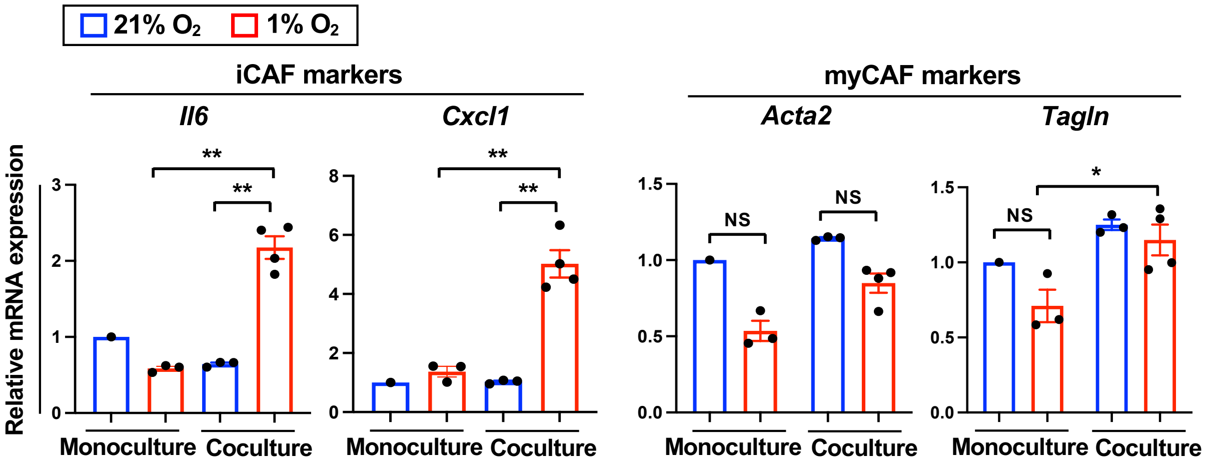


**Figure S2. Hypoxic induction of the iCAF phenotype.**

Quantitative RT-PCR analysis of iCAF and myCAF markers in CAFs (FB1) cultured alone or with mT3 tumor organoids under 21% $O_2$ or 1% $O_2$ for 72 hours (n=3-4). Expression levels were normalized by *18S rRNA*. P values were determined by two-way ANOVA with Bonferroni post-test. NS, not significant. *$p < 0.05$; **$p < 0.01$.

#### Figure S3

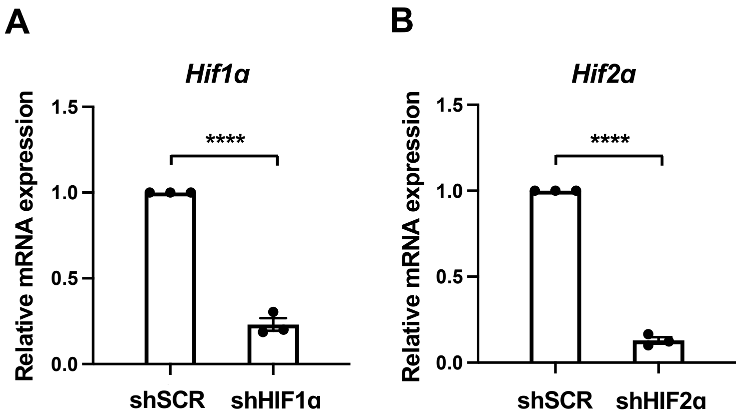


**Figure S3. Knockdown of HIF1α and HIF2α in PDAC tumor cells.**

(A) Quantitative RT-PCR analysis of *Hif1α* in mT3 tumor organoids expressing scrambled shRNA (shSCR) control or HIF1α shRNA (shHIF1α) cultured with PSCs under 1% $O_2$ for 48 hours (n=3).

(B) Quantitative RT-PCR analysis of *Hif2α* in mT3 tumor organoids expressing scrambled shRNA (shSCR) control or HIF2α shRNA (shHIF2α) cultured with PSCs under 1% $O_2$ for 48 hours (n=3).

Expression levels were normalized by *18S rRNA*. Results show mean±SEM. P values were determined by student's *t* test. ****p < 0.0001.

### Figure S4

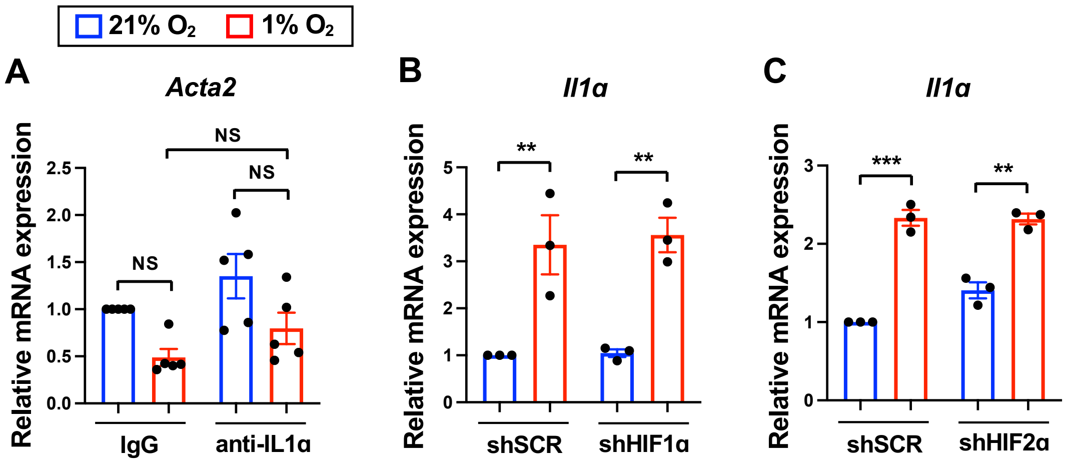


**Figure S4. IL1α does not significantly affect *Acta2* expression in PSCs and is regulated in a HIF-independent manner.**

(A) Quantitative RT-PCR analysis of *Acta2* in PSCs cultured with mT3 tumor organoids in the presence of IL1α-neutralizing antibody or isotype control antibody under 21% $O_2$ or 1% $O_2$ for 72 hours (n=5).

(B and C) Quantitative RT-PCR analysis of *Il1a* in mT3 tumor organoids expressing scrambled shRNA (shSCR) control, HIF1α shRNA (shHIF1α) (B), or HIF2α shRNA (shHIF2α) (C) cultured with PSCs under 21% $O_2$ or 1% $O_2$ for 48 hours (n=3).

Expression levels were normalized by *18S rRNA*. Each data point represents individual primary PSC lines. Results show mean±SEM. P values were determined by two-way ANOVA with Bonferroni post-test. NS, not significant. $^{**}p < 0.01$; $^{***}p < 0.001$.

**Supplemental Table S1. Primer sequences for quantitative RT-PCR. Related to Experimental Procedures.**

| Gene | Forward Primer | Reverse Primer |
|---|---|---|
| *18S rRNA* | GTAACCCGTTGAACCCCATT | CCATCCAATCGGTAGTAGCG |
| *Acta2* | TGCTGACAGAGGCACCACTGAA | CAGTTGTACGTCCAGAGGCATAG |
| *Cxcl1* | TCCAGAGCTTGAAGGTGTTGCC | AACCAAGGGAGCTTCAGGGTCA |
| *Il1α* | ACGGCTGAGTTTCAGTGAGACC | CACTCTGGTAGGTGTAAGGTGC |
| *Il6* | TACCACTTCACAAGTCGGAGGC | CTGCAAGTGCATCATCGTTGTTC |
| *Lif* | ATT GTG CCC TTA CTG CTG | TGT TAG GCG CAC ATA GCT TTT |
| *Tagln* | GCAGATGGAACAGGTGGCTCAA | CCCAAAGCCATTAGAGTCCTCTG |
